# Protein k-mers enable assembly-free microbial metapangenomics

**DOI:** 10.1101/2022.06.27.497795

**Authors:** Taylor E. Reiter, N. Tessa Pierce-Ward, Luiz Irber, Olga Borisovna Botvinnik, C. Titus Brown

**Affiliations:** Department of Population Health and Reproduction, University of California, Davis; Graduate Group in Computer Science, UC Davis; Data Sciences Platform, Chan Zuckerberg Biohub

## Abstract

An estimated 2 billion species of microbes exist on Earth with orders of magnitude more strains. Microbial pangenomes are created by aggregating all genomes of a single clade and reflect the metabolic diversity of groups of organisms. As *de novo* metagenome analysis techniques have matured and reference genome databases have expanded, metapangenome analysis has risen in popularity as a tool to organize the functional potential of organisms in relation to the environment from which those organisms were sampled. However, the reliance on assembly and binning or on reference databases often leaves substantial portions of metagenomes unanalyzed, thereby underestimating the functional potential of a community. To address this challenge, we present a method for metapangenomics that relies on amino acid k-mers (k_aa_-mers) and metagenome assembly graph queries. To enable this method, we first show that k_aa_-mers estimate pangenome characteristics and that open reading frames can be accurately predicted from short shotgun sequencing reads using the previously developed tool orpheum. These techniques enable pangenomics to be performed directly on short sequencing reads. To enable metapangenome analysis, we combine these approaches with compact de Bruijn assembly graph queries to directly generate sets of sequencing reads for a specific species from a metagenome. When applied to stool metagenomes from an individual receiving antibiotics over time, we show that these approaches identify strain fluctuations that coincide with antibiotic exposure.

## Introduction

Microbes are the most species-rich category of organisms on Earth [1], comprising an estimated 2 billion species, and yet fewer than 0.01% of species’ genomes are currently deposited in NCBI Genomes [2], their full diversity is under-described. Short read metagenomic sequencing has expanded knowledge of microbial communities and diversity [3,4,5]. In particular, metagenome assembly and annotation have produced catalogs of metagenome-assembled genomes and genes, revealing new species and functions previously unobserved in cultured organisms [3,4,6].

Along with advances in metagenome sequencing and analysis, the concept of “metapangenomics” has arisen as a framework for understanding how sets of metagenome-derived genes that are attributable to a group of organisms correlate with environmental parameters [7,8,9]. Metapangenomic methods borrow heavily from pangenome analysis. Pangenomes comprise all genomic elements – usually open reading frames or genes – found within a group of organisms and reflect the metabolic and ecological plasticity of that group [10,11]. The pangenome is divided into core and accessory genes, where core genes are shared by almost all members in the group and accessory genes are not. Core genes often encode primary metabolism or other functions necessary for a group to live in a given environment [12], while accessory genes encode functions that facilitate adaptation to changing environments [11]. The size of the pangenome (e.g. number of distinct sequences) reflects the diversity of the organisms in a pangenome (population size, number of organisms sampled) as well as the ability of those organisms to adapt to different niches [10]. Open pangenomes are those which increase indefinitely in size when adding new genomes, while closed pangenomes do not.

While pangenomes are traditionally inferred from genomes created from lab-cultured isolates (“isolates”), metapangenomics extends the ecological framework of pangenomics to metagenomes. Metapangenomics gives insight into the genes that support specific environmental adaptations by applying pangenome methods to metagenome assembled genomes (MAGs) [8], or by mapping metagenomes against isolate-inferred pangenomes [7]. Both methods give valuable insight into the presence and distribution of functional content in natural microbial communities, but either may introduce biases associated with unknown sequencing content [13]. MAGs are often incomplete or unrecoverable due to low sequencing coverage or large amounts of variation caused by SNPs, indels, rearrangements, horizontal gene transfer, sequencing error, and so on. Sequencing variation causes short read assemblers to produce unbinnable short contiguous sequences. Unbinned sequences are disproportionately comprised of genomic islands and plasmids [14], hot spots for evolution that support microbial adaptation to changing environments [15]. In contrast, read mapping against isolate-inferred pangenomes may miss functional content present in the metagenome but absent from references, especially for unknown or under-represented species.

These issues are not exclusive to metapangenome inference, and many recently developed analysis strategies overcome some of these biases. These techniques largely rely on k-mers, words of length *k* in DNA or protein sequences. Metagenome k-mer profiles contain all sequences in a metagenome, including those which may not assemble or bin, or which aren’t in reference databases. Long k-mers are also taxonomy-specific, where increasing k-mer length leads to sub-species discriminatory power [16,17]. The ability to distinguish between species without alignment or assembly have popularized the use of k-mers for metagenome analysis, primarily through lightweight sketching and compact de Bruijn assembly graphs (cDBGs). Lightweight sketching facilitates fast and accurate sequence comparisons between potentially large data sets through random but consistent sub-sampling [18,19]. cDBGs maintain connectivity between k-mers and organize them into species-specific neighborhoods [20,21].

To more fully represent the functional potential in metapangenomes, we present an analysis approach that relies on amino acid k-mers and assembly graph queries to estimate microbial (meta)pangenomes. This approach for metapangenome estimation is minimally reliant on reference databases and is assembly-free.

## Results

In an effort to reconstruct metapangenomes without loss of information from assembly and binning [14,20,21,22,23,24], we demonstrate a pipeline that relies on k-mers and assembly graphs for metapangenome estimation (**Figure 1**). We first show that amino acid k-mers (k_aa_-mers) accurately estimate microbial pangenomes by comparing amino acid profiles of proteomes (translated coding domain sequences) against the proteomes themselves (**Figure 1 A**). To derive amino acid k-mers directly from shotgun metagenome reads, we next demonstrate the accuracy of a tool called orpheum for open reading frame prediction from short sequencing reads (**Figure 1 B**). We use assembly graph genome queries to retrieve species-specific reads from the metagenome, predict open reading frames from those reads using orpheum, and build a metapangenome using protein k-mers (**Figure 1 C**). We then apply this method to species present in time-series metagenomes from a human gut microbiome.

**Figure 1:**
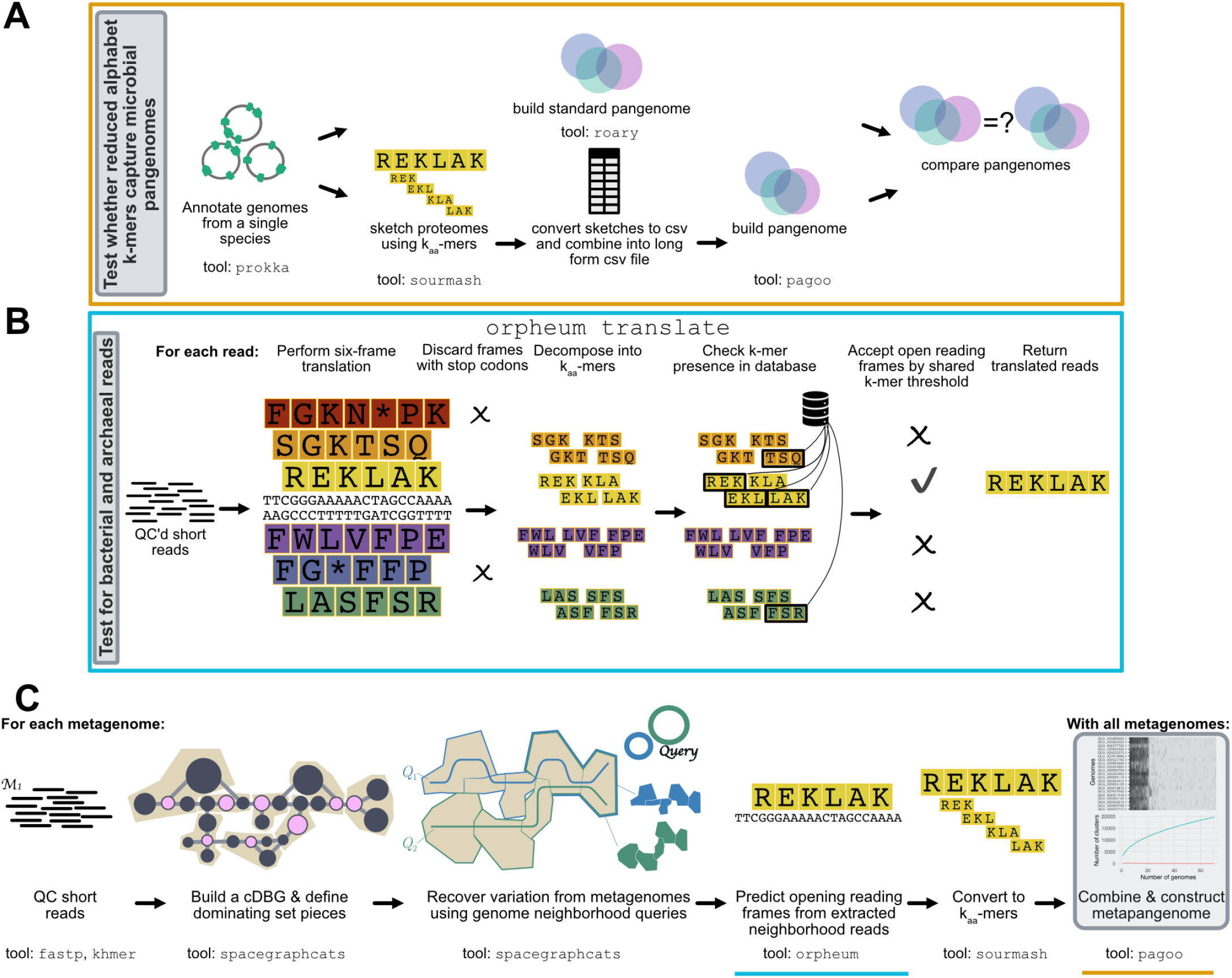
Overview of the pipeline used to build metapangenomes. Approaches that were developed or tested in this manuscript are outlined in grey. **A)** We tested whether amino acid k-mers could accurately represent bacterial and archaeal pangenomes. Using genomes annotated with prokka, we compared pangenomes built with roary, a field-standard pipeline, against pangenomes built with k_aa_-mer sketches. **B)** We tested whether open reading frames could be predicted directly from short sequencing reads using the tool orpheum. This panel is modified from [25]. **C)** We combined this approaches with metagenome assembly graph genome queries to estimate metapangenomes directly from metagenomes without assembly or binning. The blue and orange lines correspond to steps tested in panels **A** and **B**. The workflow presented in steps 1-3 of panel **C** is published in [21].

### Protein k-mers accurately estimate characteristics of microbial pangenomes

Pangenomes from isolates are typically built by assembling each isolate genome and predicting genes (open reading frames), clustering gene sequences from all genomes into a non-redundant set, and estimating the presence/absence or abundance of each gene in each genome. To determine whether bacterial and archaeal pangenomes could be constructed from protein k-mers, we compared pangenomes estimated from genes against those estimated from k-mers. We compared pangenomes from 23 species belonging to 23 phyla in the GTDB taxonomy [26], with pangenome size ranging from 20-972 genomes (mean = 203 genomes, median = 44 genomes) (**Figure S1**). For each pangenome, we computed the R^2^ between the total number of genes to the total number of k-mers, and the number of unique genes to the number of distinct k-mers within each genome. We also tested the similarity of presence/absence profiles between pangenomes constructed with different methods using the Mantel test [27].

The strength of the relationship between k-mers and genes varied dramatically for some pangenomes. Both k-mers and genes are highly correlated in DNA or protein space for most pangenomes, while a few pangenomes were outliers with low correlations (**Figure S2**). We investigated pangenomes more closely to determine the source of the poor correlations and found that they were caused by the presence of many frameshifted proteins, one of many potential criteria for exclusion of GenBank genomes from RefSeq. For example, *Leptospira interrogans* had an R^2^ of 0.12 between the total number of genes and k-mers in genomes in the pangenome, but 21 of 317 genomes contained frameshifted proteins. Removing genomes with many frameshifted proteins increased the R^2^ to 0.87 (**Figure 2 A**). This trend was consistent across pangenomes, where pangenomes with one or more frameshift-excluded genome had significantly lower R^2^ values between total number of genes and k-mers per genome than pangenomes without (Welch Two Sample t-test, estimate = −0.36, p = 0.003) (**Figure 2 B**). Other RefSeq exclusion criteria did not impact the correlation between the total genes and k-mers per genome for a given pangenome.

**Figure 2:**
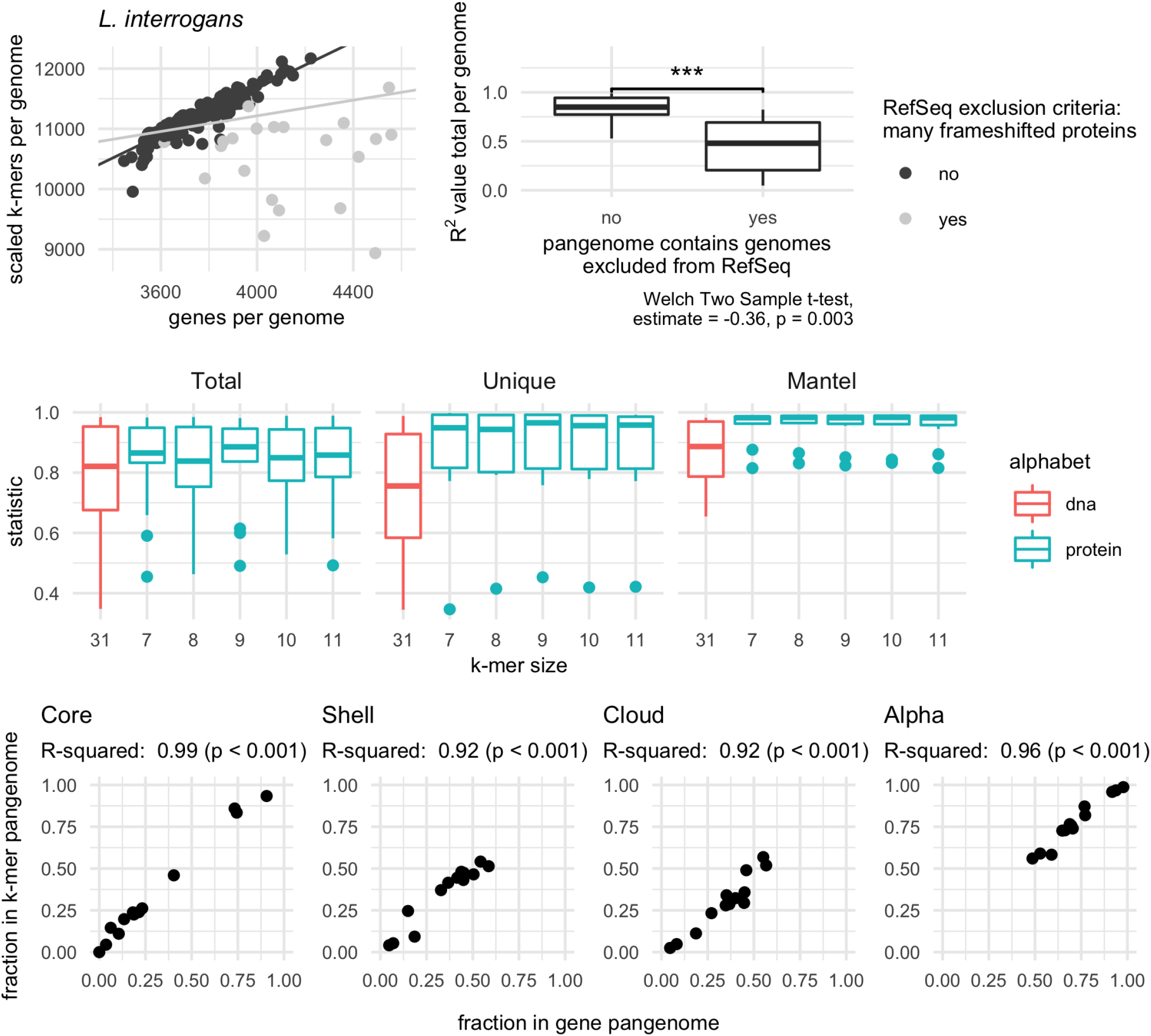
Amino acid k-mers accurately estimate characteristics of bacterial and archaeal pangenomes. **A, B)** Genomes that are excluded from RefSeq for having many frameshifted proteins reduce similarity between gene- and k-mer-based pangenomes. **A)** Scatter plot of the total number of genes and k-mers per genome for the species *Leptospira interrogans*, where each point represents a single genome in the pangenome. Removing genomes flagged with RefSeq exclusion criteria “many frameshifted proteins” improves the correlation between these variables. The light grey line corresponds to regression results when all points are used, while the dark grey line corresponds to regression results when flagged genomes are removed. **B)** Box plot of R^2^ values between the total number of genes and k-mers per genome. Pangenomes that contain genomes with the RefSeq exclusion criteria of “many frameshifted proteins” have significantly lower R^2^ values. **C)** Box plots representing the distribution of R^2^ values for linear models (Total, Unique) or statistic values for mantel tests (Mantel). Only pangenomes that do not contain genomes with the RefSeq exclusion criteria “many frameshifted proteins” are plotted. K-mer size corresponds to the number of amino acid sequences used for the k-mer for all k-mers except *k* = 31, which corresponds to the number of nucleotides. *Total* corresponds to correlations between the total number of distinct genes and k-mers in a genome. *Unique* corresponds to correlations between the number of unique genes and k-mers in genome. *Mantel* corresponds to mantel tests between the gene and k-mer presence-absence matrices. **D)** Pangenome metrics strongly correlate between gene- and k-mer-based pangenomes. Pangenome categories core, shell, and cloud refer to genes or k-mers shared between the majority (>95%), some, or singleton genomes in the pangenome. Alpha is a value from Heaps law used to estimate whether a pangenome is open or closed.

A range of k-mer sizes in amino acid alphabets had comparable performance. Using pangenomes that contained no genomes excluded from RefSeq for containing many frameshifted proteins (n = 13), we found that k-mer size had little impact on the accuracy of pangenome estimation (**Figure 2 C**). This is likely because the genomes of the same species are closely related, so protein k-mers are suffcient to overcome minor genomic variations such as those introduced by codon degeneracy or evolutionary drift [28]. The one exception was for nucleotide k-mers (*k* = 31), which did not correlate as strongly with gene-based pangenomes. This supports the use of amino acid k-mer encodings over nucleotide k-mers for construction of pangenomes. Given that neither encoding nor k-mer size impacted these performance metrics, we selected protein k-mers with *k* = 10 to complete the rest of our analysis as this combination was the only combination to fall among the top five performers across all three metrics. In addition, protein k-mers of length 10 have recently been shown to perform well for comparisons across variable taxonomic distances [17].

We next investigated whether other pangenome metrics were well correlated between our k-mer-based method and the gene-based method roary (see Methods for details). For 13 pangenomes, the percent of k-mers or genes predicted to be part of the core, shell, or cloud pangenome was strongly correlated (**Figure 2 D**). The content of the core genomes was also similar between pangenomes built with different methods. Focusing on genes or k-mers shared between all genomes in a pangenome, and limiting our inquiry to pangenomes with at least five genes shared between all genomes (n = 11), we found core k-mers contained an average of 83.9% (SD = 15.4%) of sequences in core genes, while core genes contained an average of 73.5% (SD = 16.9%) of sequences in core k-mers. This indicates congruence in the functional content represented by the core fractions of both pangenome types. Lastly, we compared whether pangenomes would be designated as open or closed by calculating the alpha value for the Heaps law model [29]. Alpha values were strongly correlated between gene- and k-mer based pangenomes (**Figure 2 D**).

Taken together, these results show that reduced alphabet k-mers can accurately estimate key characteristics of pangenomes from bacterial and archaeal genomes.

### K-mer methods accurately predict open reading frames in short sequencing reads

We next sought to determine whether open reading frames could be accurately predicted directly from short sequencing reads, as this would enable k-mer-based pangenome analysis without assembly. Without accurate open reading frame prediction, reads would need to be translated into all six translation frames prior to k-mer decomposition. This would inflate the number of k-mers and decrease similarity between genomes.

We evaluated whether orpheum, a tool recently developed to predict open reading frames in Eukaryotic short reads [25], could also perform this task in bacterial and archaeal sequences. Orpheum predicts open reading frames by comparing reduced alphabet k-mers in six frame translations of short sequencing reads against those in a database (containment) and assigns an open reading frame as coding if containment exceeds a user-defined threshold [25]. To evaluate orpheum, we constructed a database containing all k-mers in coding domain sequences from genomes in GTDB rs202. Using representative genomes from the 23 species above, as well as 20 additional RefSeq genomes not in the GTDB rs202 database, we simulated short sequencing reads either from coding domain sequences or non-coding sequences and used these reads to test orpheum.

Using default parameters, orpheum accurately separated coding from non-coding reads when reads were simulated from genomes in GTDB (**Figure 3 A**). On average, 5.3% (SD = 2.8%) of reads that were coding were incorrectly predicted to be non-coding, while 4.9% (SD = 1.5%) of reads that were non-coding were incorrectly predicted to be coding. For reads simulated from genomes not in GTDB, orpheum recovered the majority of coding reads when genomes of the same species were in the database (**Figure 3 A,B**). On average, 30.2% (SD = 27.1%) of reads that were coding were predicted to be non-coding, while 4.8% (SD = 5.5%) of reads that were non-coding were predicted to be coding. Accuracy decreased with increasing taxonomic distance between the query genome and the closest relative in the database (**Figure 3 B**).

**Figure 3:**
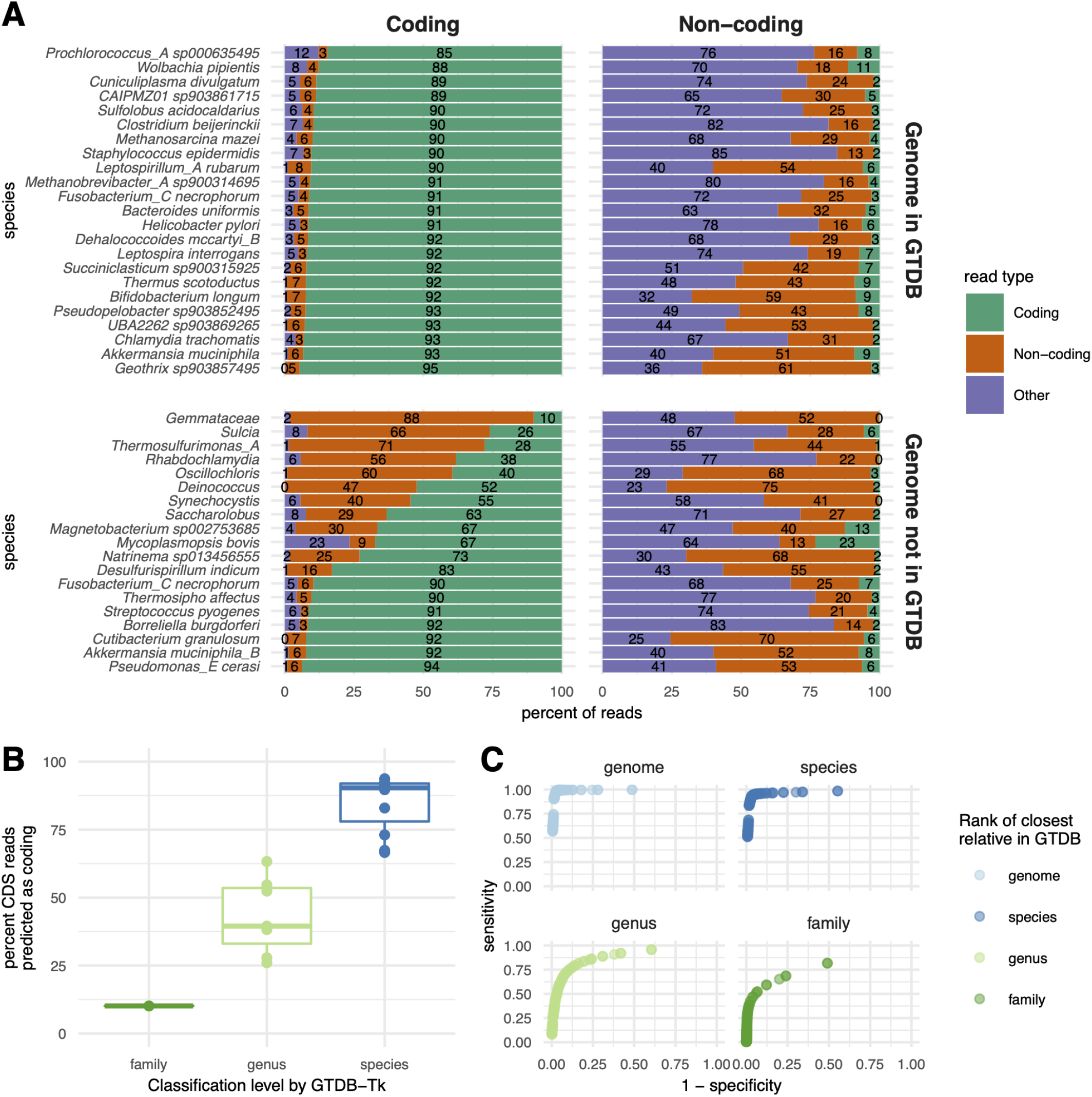
Orpheum correctly assigned short sequencing reads as coding or non-coding and selects the correct open reading frame. **A)** Percent of simulated coding or non-coding sequences predicted as coding, non-coding, or discarded based on quality metrics (see methods). Genomes are split by those in GTDB and those not in GTDB. Genomes not in GTDB are labelled by taxonomic assignment from GTDB-tk. Predictions were made using default parameters (Jaccard containment = 0.5). **B)** Boxplots of the percent of coding reads that were recovered by Orpheum, separated by the level of taxonomic assignment achieved by GTDB-Tk. Orpheum recovers more coding sequences when there are closely related genomes in the database. **C)** Receiver operating curves for the Jaccard containment thresholds. Curves are separated by the level of taxonomic assignment achieved by GTDB-Tk, and values are averaged across all genomes that fell within those categories. The best Jaccard threshold decreases when there are fewer closely related genomes in the database.

For genomes that had at least species-level representatives in GTDB, the largest source of error was non-coding reads being predicted as coding (**Figure 3 A**). We hypothesized that these reads originated from pseudogenes as these sequences would likely not be annotated as coding in the genomes from which the reads were simulated from, but may retain some k-mers contained in the database. To assess this hypothesis, we used annotation files produced by the NCBI Prokaryotic Genome Annotation Pipeline (PGAP), which annotates pseudogenes, for the 23 genomes for which these files were available [30,31]. On average, 12.4% (SD = 13.8%) of non-coding reads that were predicted to be coding fell within pseudogenes annotated by the PGAP pipeline. We then BLASTed a subset of the remaining non-coding reads that were predicted to be coding against the NCBI nr database. All reads we investigated had at least one match at 100% identity to protein sequences in the database, suggesting our test genomes contained additional pseudogenes not annotated by PGAP, or that the software we used to predict open reading frames missed some coding sequences (see Methods). Because this method of open reading frame prediction cannot distinguish pseudogenes, it may not be appropriate for species with many pseudogenes.

Some coding sequences were also predicted to be non-coding. We hypothesized that this was caused by sequencing error introduced into the simulated reads. We mapped the simulated reads against the coding domain sequences from which they were derived and calculated mapping error rates. While all reads mapped, the error rate was higher for reads that were predicted to be non-coding than those predicted to be coding (Welch Two Sample t-test, estimate = 0.00523, p < 0.001).

Protein k-mers from predicted open reading frames in the simulated short sequencing reads recapitulated similarity between genomic coding domain sequences. We estimated the Jaccard similarity between genomes using k_aa_-mers (*k* = 10) from annotated coding domain sequences, and compared this against Jaccard similarity between genomes using k_aa_-mers from predicted open read frames in the simulated short sequencing reads. Genomes that were most similar in one matrix were also most similar in another matrix (Mantel statistic = 0.9975, p < 0.001). The average similarity among all pairwise comparisons for the coding domain sequences was 2.6%, and this decreased to 2.5% when using the open reading frames predicted from reads. This demonstrates that information recovered from open reading frame prediction from short read is similar to that derived directly from the genome sequence.

The majority of predictive capability originated from species-level databases. We performed ORF prediction using just species-level databases for genomes that had at least a species-level representative in GTDB, and compared this against ORF prediction using the full GTDB database. On average, there was no change between the percent of reads derived from coding domain sequences when a species-level database was used versus when all of GTDB was used to predict open reading frames (**Figure S3**).

Decreasing the Jaccard containment threshold increased the sensitivity and specificity of ORF prediction when there are no closely related genomes in the database (**Figure 3 C, Table 1**). The Jaccard containment threshold controls the final prediction of coding vs. non-coding, as well as the the number of open reading frames which a read is translated into. On average, increasing the rank of the closest taxonomic relative in the database by one taxonomic level decreased the optimal Jaccard containment threshold by 0.13. We note that orpheum performed well, achieving sensitivity > 0.9 or better, when genomes of the same strain, species, and genus are present, but decreases significantly when the next closest relative is at the family level (**Figure 3 C**).

**Table 1:**
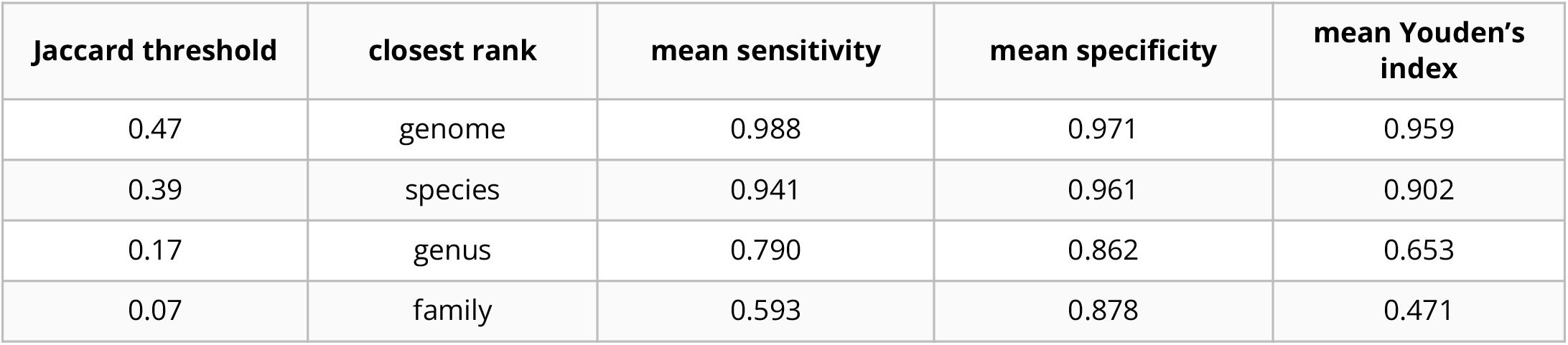
Jaccard containment thresholds that maximize the Youden’s index depending on the taxonomic rank of the closest relative in GTDB.

| Jaccard threshold | closest rank | mean sensitivity | mean specificity | mean Youden's index |
| --- | --- | --- | --- | --- |
| 0.47 | genome | 0.988 | 0.971 | 0.959 |
| 0.39 | species | 0.941 | 0.961 | 0.902 |
| 0.17 | genus | 0.790 | 0.862 | 0.653 |
| 0.07 | family | 0.593 | 0.878 | 0.471 |

Overall, these results show that open reading frames can be accurately determined from short sequencing reads when closely related proteomes are available.

### K-mer-based metapangenomics combined with assembly graphs reveals strain dynamics

Given that amino acid k-mers accurately estimated pangenomes, and that the correct open reading frame could be predicted reliably from short sequencing data, we next combined these approaches to perform metapangenome analysis from short read shotgun metagenomes. We used 12 metagenomes from a single individual sampled over the course of a year by the Integrated Human Microbiome Project (iHMP) [32]. The individual was diagnosed with Crohn’s disease, a sub type of inflammatory bowel disease characterized by inflammation along the gastrointestinal tract. The individual received three courses of antibiotics over the year and each course was separated by weeks without antibiotics (**Figure 4 A**).

**Figure 4:**
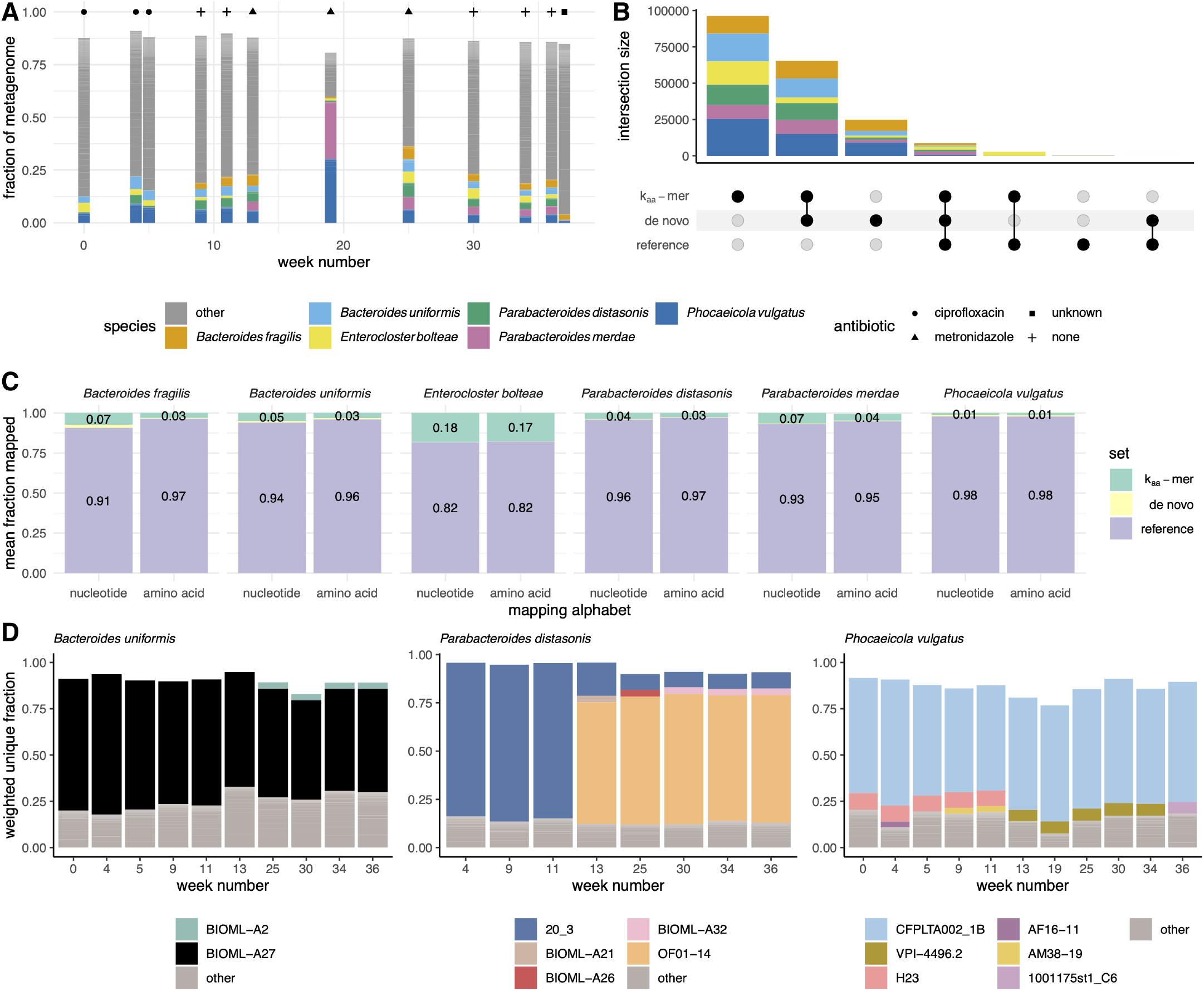
K_aa_-mer metapangenomes reveal species and strain dynamics in time series gut microbiome metagenomes after antibiotic exposure. **A)** Antibiotic courses and corresponding gut microbiome profiles for a single individual with Crohn’s disease. Fractional abundances are colored by species, with only the six species that accounted for greater than 2% of all metagenome reads displayed. **B)** Upset plot of amino acid k-mers (*k* = 10) present in the k_aa_-mer metapangenomes, the *de novo* metapangenomes, and the reference pangenome. Intersections are colored by species. **C)** Bar plots indicating the average fraction of reads used to build the k_aa_-mer metapangenome that mapped first against the reference pangenome, then against the *de novo* metapangenome, or were unmapped. More reads mapped in amino acid space than in nucleotide space. Only the fraction of reads that mapped to the reference pangenome and the fraction reads that were unmapped are labelled. **D)** Bar plots of the fraction of k_aa_-mer metapangenome sequences that were anchored to a given strain using the sourmash gather algorithm against the GTDB rs202 database (*k* = 51). Colors represent strains, which are labelled by their NCBI strain name. Missing fractions depicted as blank space between the bar and one represent novel k-mers not in the database. Only genomes that accounted for greater than 2% of the weighted fractional abundance and that were annotated as the same species are colored. Weeks in which the species was low-abundance are excluded. Starting at at week 13, sequences from previously unobserved strains were detected within each metapangenome. This timing coincides with metronidazole administration.

We estimated the metapangenome for each species that was detected in all 12 metagenomes and that accounted for at least 2% of reads across metagenomes, for a total of six metapangenomes (**Figure 4 A**). To obtain all sequencing reads that originated from genomes of these species, we performed assembly graph genome queries [21]. Assembly graphs contain all sequences in a metagenome, and assembly graph queries return sequences in the metagenome that are either in the query or nearby to the query in the graph. Assembly graph genome queries return sequencing reads that originate from genomes in the metagenome that have as little as 0.1 Jaccard similarity (approximately 93% average nucleotide identity (ANI) [17]) to the query genome [21]. After retrieving reads in this way, we predicted open reading frames using orpheum. We used species-level databases as these were successful in the context of isolate genomes not in the database (see above) and because they would be more likely to filter out reads beyond the species boundary (95% ANI [33]) that were returned by assembly graph queries. Using the predicted amino acid sequences, we built k_aa_-mer metapangenomes for each of the six species (**Figure 1 C, Figure S4, Table S1**).

We compared these metapangenomes against reference pangenomes built using genomes of the same species in GTDB and against *de novo* metapangenomes built from MAGs of the same species that were assembled and binned from these samples (see Methods). Almost all sequences from the reference pangenome occurred within the k_aa_-mer metapangenomes (**Figure 4 B**), indicating we recovered the majority of sequencing variation contained within the reference pangenome. Further, a large fraction of sequences were shared between the *de novo* metapangenome and the k_aa_-mer metapangenome (**Figure 4 B**), indicating we also recover the majority of variation captured by assembly and binning.

A large fraction of k-mers were only represented in the k_aa_-mer metapangenome (**Figure 4 B**). To determine whether these sequences represented true biological variation from our query species and not contamination from other species, we next iteratively mapped the reads that were used to build the k_aa_-mer metapangenome against the reference pangenome and the *de novo* metapangenome (**Figure 4 C**). The majority of reads mapped against the reference pangenomes (mean = 92.2%, SD = 12.4%), a few of the unmapped reads mapped against the *de novo* metapangenome (mean = 0.8%, SD = 1.3%), and 7.1% (SD = 12.3%) of reads did not map. We repeated this process a second time but mapped in amino acid space. Mapping in amino acid space improves sensitivity over nucleotide mapping [34]. The fraction of reads that mapped increased by an average of 1.7% (Welch Two Sample Paired t-test, estimate = 1.9, p < 0.001), accounting for 94.6% of total reads and indicating that a substantial fraction of distinct sequences in the k_aa_-mer metapangenome represent diverged sequences with similar amino acid sequences.

On average, 5.4% (SD = 12%) of reads from the k_aa_-mer metapangenome were unaccounted for after mapping against other (meta)pangenomes. We assembled and annotated these reads, and BLASTed the 88 resultant protein sequences against the NCBI nr database. Of 56 predicted genes with a BLAST hit, 76.8% matched sequences from the same species or closest specified lineage level as the top hit. This suggests our method recovers functional sequences even if those sequences are not in MAGs or in reference databases (in this case, in NCBI nr but not GTDB).

Visualizing the k_aa_-mer metapangenomes alongside sequencing depth information, we observed dynamics in the presence of species (**Figure S4 A**) or strains (**Figure S4 B**) in response to antibiotic administration. The fluctuation of the presence of species indicates that antibiotic administration impacted the community structure of the gut microbiome, as is expected [35]. This is exemplified by periodic blooms of *Enterocloster bolteae*, an organism associated with disturbance succession [36].

Similarly, we detected changes in accessory k_aa_-mers, the majority of which appeared or disappeared on or after the start of metronidazole administration at week 13 (**Figure S4 B**). Metronidazole targets anaerobic bacteria via reduction by pyruvate:ferredoxin oxidoreductase system which creates an electron sink that produces free radicals that are toxic to cells [37]. Metronidazole treatment disproportionately impacts the presence of anaerobes in the gut microbiome [38]. We hypothesized that fluctuations in accessory k_aa_-mers reflected strain-level turn over in the community. To confirm this, we compared the nucleotide k-mer content in each query neighborhood against the GTDB database to determine which strains were present (**Figure 4 D**). We used a k-mer size of 51, as this is indicative of strain-level similarity [16,17]. In each of the three species we investigated, we identified different patterns of strain fluctuations. In *Bacteroides uniformis*, only one strain of *B. uniformis* (BIOML-A27) was present until week 25, when another strain appears (BIOML-A2). In *Parabacteroides distasonis*, the dominant strain switches at week 13 (20_3 to OF01-14), coincident with other strains appearing (BIOML-A21, BIOML-A26, BIOML-A32). Lastly, in *Phocaeicola vulgatus*, the dominant strain does not change through the time course, but multiple other strains are detected with one strain switch occurring at week 13 (H23 and AM38-19 to VPI-4496.2).

Taken together, these results demonstrate that k_aa_-mer metapangenomes built from metagenome assembly graph neighborhoods can capture species and strain dynamics in microbiome communities.

## Discussion

We present a method to perform assembly-free metapangenomics that is minimally reliant on reference databases. We show that pangenome metrics like core, cloud, and shell pangenome fractions can be accurately estimated with amino acid k-mers (*k* = 10). We then demonstrate accurate prediction of open reading frames in highly accurate short sequencing reads by comparing amino acid k-mers in all translation frames against a database of k-mers from bacterial and archaeal genomes in GTDB (rs202). Combining these tools enables pangenome estimation directly from quality controlled short sequencing reads. In the context of metagenomes, these approaches enable metapangenome estimation without the need to *de novo* assemble and bin sequences, eliminating common sources of lost sequencing variation [21]. These techniques also reduce the dependence of metapangenomics on complete or comprehensive reference databases, which can be important for understudied environments.

While we leverage open reading frame prediction from short reads and k_aa_-mer pangenomes in the context of metapangenomes, these approaches have additional applications. Open reading frame prediction with orpheum can be executed on microbial Illumina short read data sets. This may improve functional recall from metagenome short reads from organisms without reference genomes and from communities that have low assembly rates [39], or from metatranscriptomes, however these applications must be validated. Similarly, k_aa_-mer pangenomes built from sketches may offer benefits over other pangenome construction methods, many of which are predominately tied to exact matching of k-mers between genomes. Exact matching of k-mers between genomes enables additional genomes to be added to the pangenome without having to re-cluster gene sequences. Exact matching also allows direct comparisons to distantly related organisms, which creates a unified framework for genome comparisons even when organisms are distantly related. While we predominantly explored k_aa_-mers of length 10 in this paper, k-mers from other degenerate alphabets like Dayhoff or hydrophobic-polar encodings may allow for accurate comparisons at greater evolutionary distances [17,25]. When pangenomes are built from FracMinHashes as performed here, the scaled down sampling parameter may also enable faster pangenome size estimation even for very large collections of genomes and could be potentially used a quality control metric for annotated genomes. While developed for the metapangenomics space, this study demonstrates that k_aa_-mer pangenomes also accurately estimate pangenomes built from isolate genomes. Since building k_aa_-mer sketches and searching for exact matches of k_aa_-mers between genomes is fast, this provides an alternative approach for building pangenomes. Lastly, our results suggest that the number of k_aa_-mers in a genome strongly correlates with the number of genes; we observed that while the number of genes per genome is increased for genomes with the RefSeq exclusion criteria of “many frameshifted proteins”, there is no commensurate increase in the number of k_aa_-mers observed. This suggests that the number of k_aa_-mers in a genome could be used to predict the expected range of predicted genes, and could be used as a quality diagnostic criteria.

We posit that the combination of these approaches is potentially most useful in the context of analyzing metagenome assembly graphs. Assembly graphs like compact de Bruijn graphs (cDBG) capture all sequences in a metagenome, including sequences with high strain variation or low coverage, which may not be captured by other analysis methods. A targeted query of an assembly graph, for example with a metagenome-assembled genome bin, can recover all sequencing reads in a metagenome that originate from all genomes of the same species [21]. Often, reads may be longer than unitigs (nodes) in the cDBG in regions of high variation making read retrieval desirable. While recovering these reads and assigning their taxonomic identity through graph queries is useful, many of the recovered reads cannot be assembled due to prolific sequencing variation attributable to strain diversity in the original microbial community. Yet, the sequences represented by these un-assembleable reads often encode functional potential, some of which may be key to a microorganisms functioning within its ecosystem [20,40]. The approaches presented in this paper enable these sequences to be represented in metapangenome estimation. Indeed, as demonstrated by the number of bins generated per species at each time point, many species were observed in a sample for which no bin was produced, often because the species was present at low abundance (**Figure S4**). K_aa_-mer metapangenomes rescue these sequences and include them in the analysis.

Interestingly, in some samples, *de novo* metagenome analysis produced multiple MAGs (**Figure S4**). While not explored here, it may be possible to estimate the number of strains present in a sample using k_aa_-mer abundances. FracMinHash sketches optionally retain abundance information, so it is possible that this information could be retrieved directly from these sketches.

This method is not without shortcomings, the largest of which is the current lack of integration of k_aa_-mers with functional annotations. We see three avenues by which this could be ameliorated. First, the underlying cDBG used to produce assembly graph query neighborhoods could be annotated, and these annotations could be paired with k_aa_-mers. The technology to do this exists [41,42], however this approach is the least scalable option. Second, using the sequences included in the orpheum species databases, species-level protein databases that include functional annotations could be built and searched using sourmash gather. While rapid, this approach is undesirable given its reliance on existing annotations for genes within a given species. Lastly, a database such as the NCBI nr database could be made searchable in a similar capacity, increasing the breadth of possible matches. This may require new algorithm development to maintain scalability [43].

While cDBGs are often common intermediary data structures in bioinformatics pipelines (e.g., metagenome assembly), the organization of cDBGs is still under explored. Sequences from a genome or from closely related genomes often co-locate in neighborhoods of the cDBG [20,21], but regions of high sequence conservation and horizontally transferred genes create bridges between otherwise distant neighborhoods in the graph [20]. Even still, species-level neighborhoods are well conserved and can be extracted [21,44]. As we observed in strain-level plots of query neighborhoods wherein a fraction of the graph fell in the “other category” and belonged to low-abundance strains of the same species or organisms of other closely-related species (**Figure 4 D**), it is unclear the extent to which assembly graph organization breaks down in real metagenomes. Prolific horizontal gene transfer could account for shared sequencing content, either by collapsing neighborhoods in the graph or by increasing the fraction of shared sequences between community members. Deeply sequenced paired long and short reads from the same community could help resolve these questions.

While we observe strain dynamics, the ecological reason for those signals cannot be determined from short read sequences alone; the patterns we observe could be the result of strain turn over, the presence of multiple strains, horizontal gene transfer between strains, or some combination of all of these influences. Again, long read sequencing, or sequencing from isolate genomes, could resolve these questions, particularly as long read lineage-resolved methods become mainstream [45]. In principle, the methods presented here should extend directly to long read sequences, however this remains a point of future research.

## Methods

All code is available at github.com/dib-lab/2021-panmers (results section 1; DOI: 10.5281/zenodo.6761161), github.com/dib-lab/2021-orpheum-sim (results section 2; DOI: 10.5281/zenodo.6761169), and https://github.com/dib-lab/2021-metapangenome-example (results section 3; DOI: 10.5281/zenodo.6761180).

### Selection of benchmarking species for pangenome analysis

We selected a species representative for each of the 23 phyla in GTDB rs202 [26]. To select representative species, we first removed species with fewer than 20 genomes and greater than 1000 genomes. While this approach scales beyond 1000 genomes, we elected to benchmark smaller sets to iterate over the potential parameter space more quickly. Of species remaining after filtering, we selected the species within each phyla that had the largest number of genomes. We downloaded these genomes from GenBank. Species names are recorded in **Figure S1**.

### Calculating the gene-based pangenome with roary

To calculate the gene-based pangenome, we first annotated each genome using prokka [46]. We then used the resulting GFF annotations files to calculate the pangenome with roary using default settings [47].

### Calculating the k-mer based pangenome with sourmash

To calculate k-mer based pangenomes, we used sourmash `sketch` to generate signatures from the prokka-predicted amino acid sequences (`.faa` files) [48]. We used the protein alphabet (*k* = 7, 8, 9, 10, 11), dayhoff alphabet (*k* = 13, 15, 17), and the hydrophobic-polar alphabet (*k* = 27, 31). All signatures were calculated with a scaled value of 100. The scaled parameter controls the fraction of the total k-mers represented by the sketch; a scaled value of 100 indicates that 1/100th of the distinct k-mers in a genome were included in each sketch. We converted signatures from json format into a genome x hash presence-absence matrix.

### Correlating gene-based and k-mer based pangenomes

Using the presence-absence matrices for the gene-based and k-mer-based pangenomes, we correlated total genes/k-mers observed per genome and total unique genes/k-mers observed per genome for each species. We used the `rowSums()` function in R to determine the number of genes/unique genes per matrix, then used the `lm()` function with default parameters to correlate the values. We also used the Mantel test to determine whether genomes that were most similar in the gene presence-absence matrix were also most similar in the k-mer presence-absence matrix [27]. We used the `mantel()` function in the R vegan package to perform this test [49]. We used distance matrices calculated with the `dist()` the mantel test. function using the parameter `method = “binary”` as input to

To estimate the containment between the sequences in core genomes as estimated by roary and as estimated by k_aa_-mers, we limited the core genome to sequences present in all genomes. We then generated FracMinHash protein sketches using sourmash `sketch` (*k*=10, scaled=100) for roary core sequences and k_aa_-mer core sequences and estimated containment using sourmash `compare` [48].

### Generating standard pangenome metrics with pagoo

The pagoo R package provides functions to analyze bacterial pangenomes [50]. We used this package to generate standard pangenome metrics and visualizations. These metrics are based on the presence-absence matrices generated above and include calculation of the core, shell, and cloud genome sizes and estimation of the alpha value in Heaps law for estimation of pangenome openness.

### Augmenting benchmarking species set to include genomes not in GTDB for open reading frame prediction

We next generated a benchmarking data set for open reading frame prediction. We selected a genome from each of the 23 species evaluated above, choosing the GTDB rs202 representative genome for each species. Genome accessions are recorded in **Table S2**. Given that open reading frame prediction relies on a database, and we used k-mers in GTDB rs202 to generate this database, we also wanted to select genomes that were not in GTDB to evaluate this method. We determined the bacterial and archaeal genomes that were added to RefSeq after the construction of GTDB rs202 (April 2021-November 2021). From this set, we selected a representative genome from each of the distinct NCBI phyla represented among these genomes, 20 in total. Genome accessions are recorded in **Table S3**. We then ran GTDB-tk on these genomes to predict the GTDB taxonomy of each [51].

### Simulating coding domain sequence and non coding domain sequence reads with polyester

We next created a labelled data set of simulated reads that were generated from either coding domain sequences (CDS) or non-coding regions within each genome. We annotated the genomes with bakta to produce CDS ranges [52], and used polyester to simulate reads from CDS or non-coding regions [53]. We used the default short read error profile within polyester.

### Determining short read open reading frames with orpheum

We used the orpheum tool to predict open reading frames from simulated short reads [25]. Orpheum was developed to predict open reading frames in short RNA-seq reads from Eukaryotic organisms without a reference genome or transcriptome sequence [25]. Orpheum performs six-frame translation on nucleotide sequencing reads, calculates k-mers in an amino acid encoding at the designated k-mer length, and then estimates the containment of k-mers in a reference database in each translated frame. It then selects all open reading frames based on a containment threshold, and returns those reads as translated amino acid sequences. Open reading frames are excluded if they contain stop codons, low complexity sequences, or if the read is too short to perform translation.

Reads are designated as non-coding if they don’t reach the containment threshold and are not excluded for other reasons. We constructed a database from all genomes in GTDB rs202. We either downloaded predicted open reading frames from GenBank, or generated them using prodigal [54], and translated them into protein sequences using transeq [55]. We then used to build a nodegraph using a k_aa_-mer size of 10.

### K_aa_-mer metapangenome analysis of iHMP metagenomes

We used sourmash, spacegraphcats, and orpheum to perform k_aa_-mer metapangenome analysis of 12 iHMP time series gut microbiomes captured by short read shotgun metagenomes [56]. We downloaded samples HSM6XRQB, HSM6XRQI, HSM6XRQK, HSM6XRQM, HSM6XRQO, HSM67VF9, HSM67VFD, HSM67VFJ, HSM7CYY7, HSM7CYYD, HSM7CYY9, HSM7CYYB from ibdmdb.org. We adapter and quality trimmed each sample with fastp (parameters `--detect_adapter_for_pe`, `qualified_quality_phred 4`, `--length_required 31`, and `--correction`) [57], removed human host sequencing reads with bbduk (parameters `k=31`, reference file https://drive.google.com/file/d/0B3llHR93L14wd0pSSnFULUlhcUk/edit?usp=sharing), and k-mer trimmed reads using khmer `trim-low-abund.py` (parameters `-C 3`, `-Z 18`, `-V`) [58]. We then `gather` used sourmash to infer the taxonomic profile of each sample, using the GTDB rs202 database (*k* = 31, https://osf.io/w4bcm/) [56]. We summarized the results to species-level using the GTDB taxonomy. We retained species with a cumulative sum of at least 2% (sum of `f_unique_to_query`) across metagenome reads as query genomes. We downloaded each genome from GenBank (**Table 2**) and performed spacegraphcats assembly graph queries with each (parameters `ksize: 31`, `radius: 1`, `paired_reads: true`) [21]. Using the returned reads, we predicted open reading frames using orpheum `translate` (parameters `--jaccard-threshold 0.39`, `--alphabet protein`, `--peptide-ksize 10`) and using species-level GTDB databases. We sketched each set of translated reads using sourmash `sketch` (parameters `protein, -P k=10,scaled=100,protein`) [48], converted each sketch to a csv file, and then combined csv files for a single query species across all metagenomes. This long format csv was used as input for the R pangenome package pagoo, using the `pagoo()` function [50]. We used pagoo methods `pg$gg_binmap()`, `pg$summary_stats()`, and `pg$pg_power_law_fit()` to visualize the pangenome, calculate the size of the core, shell, and cloud, and estimate alpha.

**Table 2:** Query genome GTDB species names and GenBank accessions.

| species | accession |
| --- | --- |
| *Parabacteroides distasonis* | GCA_000162535.1 |
| *Enterocloster bolteae* | GCF_003433765.1 |
| *Bacteroides fragilis* | GCF_003458955.1 |
| *Parabacteroides merdae* | GCF_003475305.1 |
| *Bacteroides uniformis* | GCF_009020325.1 |
| *Phocaeicola vulgatus* | GCF_009025805.1 |

### Comparing k_aa_-mer, *de novo*, and reference (meta)pangenomes

We compared the k_aa_-mer metapangenomes against other (meta)pangenomes. We first constructed a reference pangenome for each query species as detailed in the methods section, “Calculated the gene-based pangenome with roary.” We used all genomes in the GTDB rs202 database for a given species (**Table 2**). We then constructed *de novo* metapangenomes for each species. Using quality controlled reads from each sample, we assembled each metagenome separately using megahit with default parameters [59]. We binned the resultant assemblies using metabat2 with parameter `–m 1500` [60]. We assigned GTDB species to each bin using sourmash `gather` (DNA, *k* = 31, scaled = 2000) against the GTDB rs202 database, selecting the species of the best match [19]. We decontaminated each bin with charcoal using default parameters [61]. We annotated each bin using prokka [46]. To compare the sequence content of the (meta) pangenomes, we sketched the protein sequences from the *de novo* and reference (meta) pangenomes using sourmash `sketch` (protein, *k* = 10, scaled = 100). We intersected the hashes in these sketches to assess shared sequencing content, and visualized it using the R complexUpset package [62].

To further compare the sequence content of the k_aa_-mer metapangenomes to the *de novo* and reference (meta)pangenomes, we mapped the reads used to build the k_aa_-mer metapangenomes iteratively against the (meta)pangenomes. We first mapped the reads against the reference pangenome using bwa `mem` with default parameters [63]. We used samtools `stat` to determine read mapping statistics, and samtools `view` and `fastq` to extract the unmapped reads [64]. We then mapped the unmapped reads against the *de novo* metapangenome. To prepare the *de novo* metapangenome for mapping, we clustered nucleotide sequences at 95% using cd-hit-est [65]. We then mapped the unmapped reads using bwa `mem` with default parameters. We extracted unmapped reads using the same procedure as above. To determine whether more reads mapped in amino acid space, we translated the reference and *de novo* (meta)pangenomes into protein sequences using transeq [55], and then repeated the same iterative mapping procedure using paladin `align` [34]. To test whether the fraction of mapped reads increased between the two mapping protocols, we used the R function `t.test()` using parameter `paired = TRUE`.

## Appendix/Supplementary information

**Figure S1:**
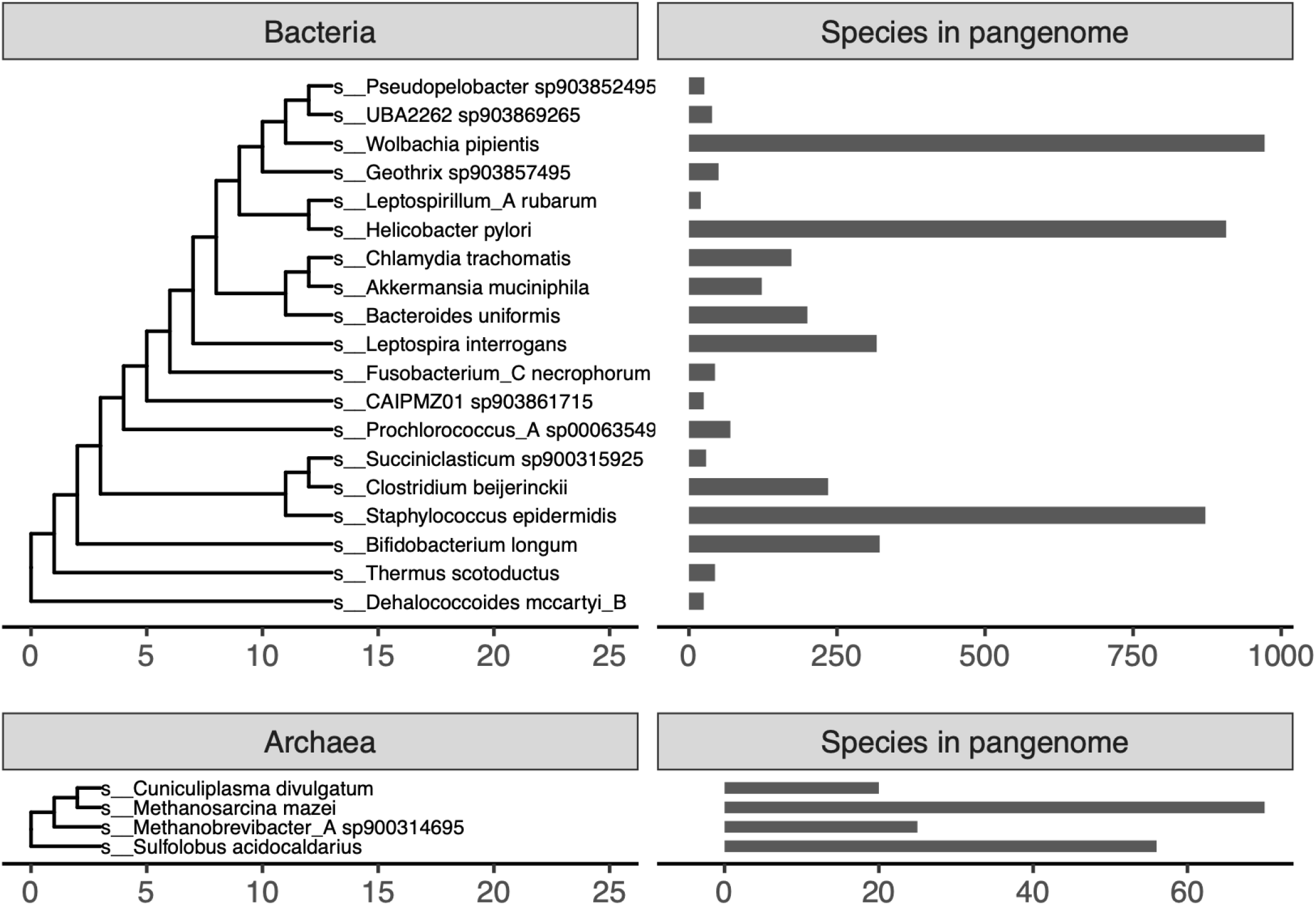
GTDB species used in this paper. These species were used to benchmark pangenome construction with reduced alphabet k-mers and open reading frame prediction from short sequencing reads. The trees are the default GTDB rs202 trees, with tips representing species not used in this paper removed.

**Figure S2:**
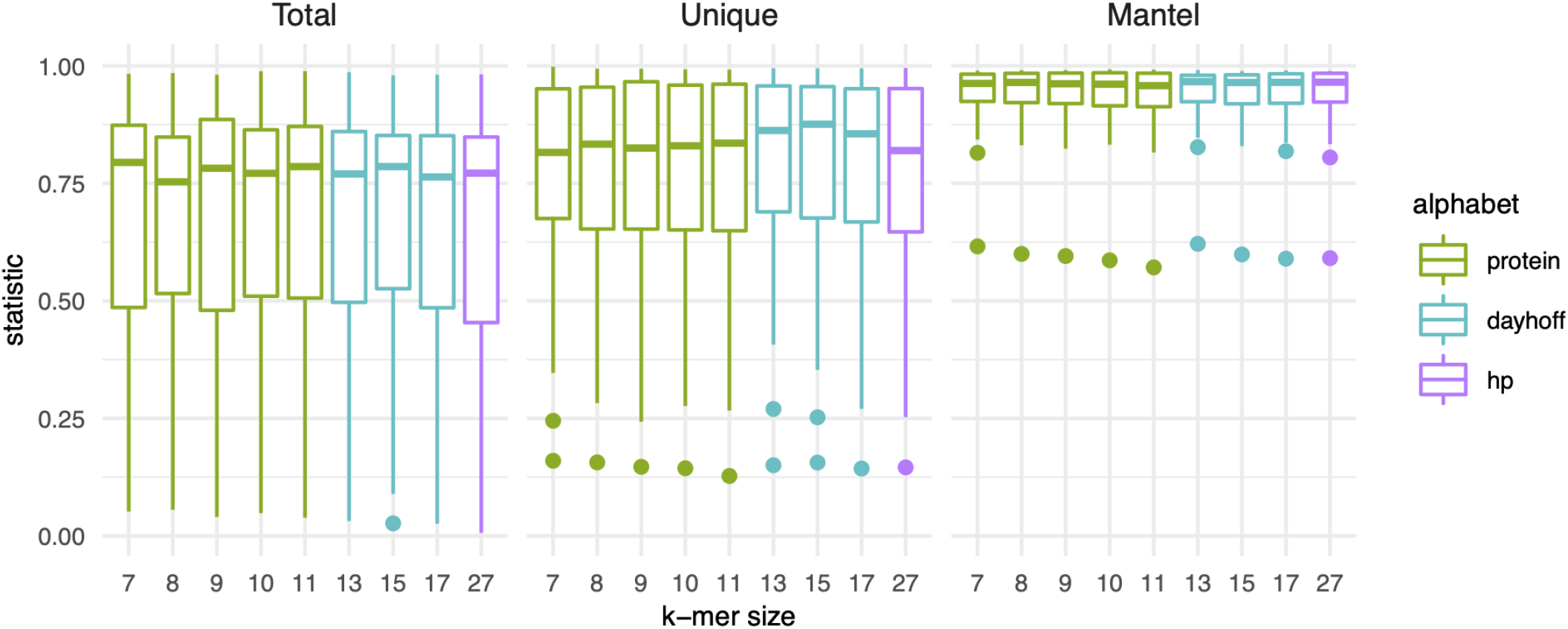
K-mer size and encoding do not impact pangenome estimation with k-mers. Box plots representing the distribution of R^2^ values for linear models (Total, Unique) or statistic values for mantel tests (Mantel) calculated for each pangenome. All pangenomes are included, whether they contain genomes with the RefSeq exclusion criteria “many frameshifted proteins” or not. See figure legend for **Figure 2** for a description of Total, Unique, and Mantel.

**Figure S3:**
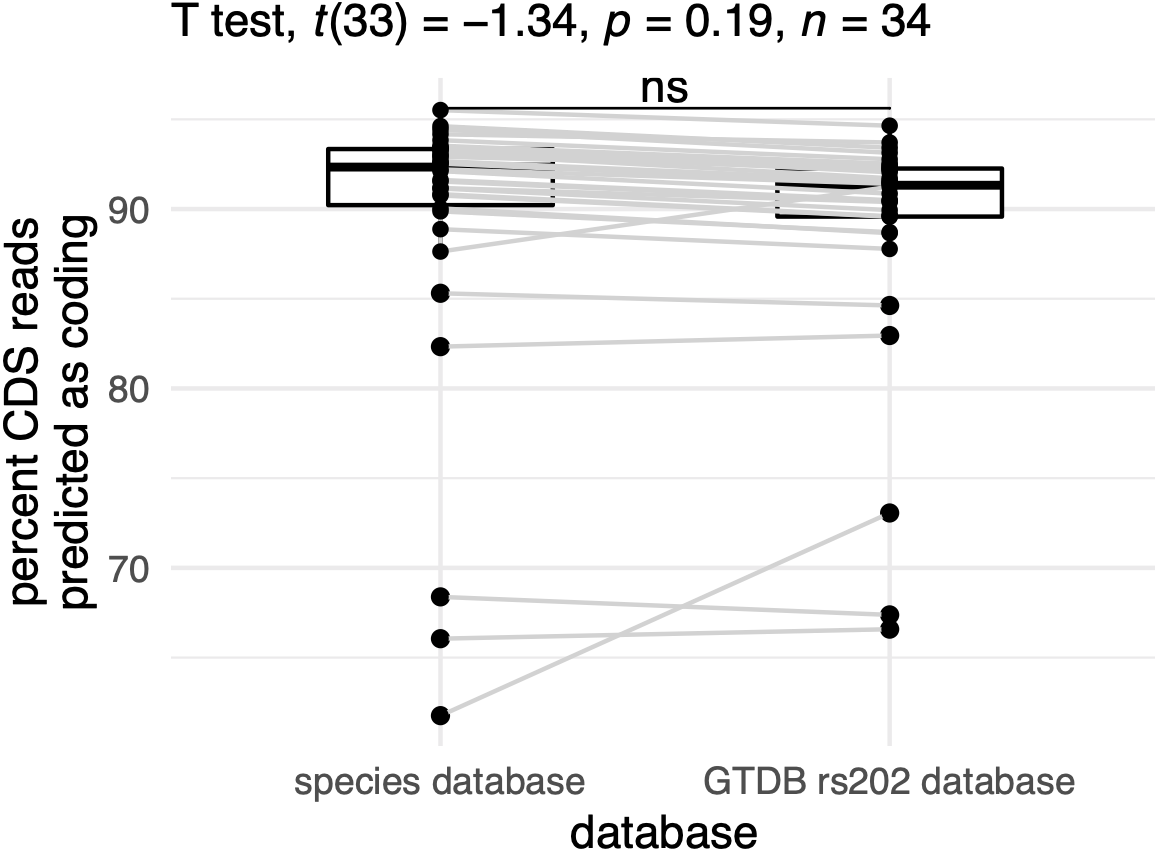
Percent of reads encoding coding domain sequences (CDS) that were predicted to be coding. There is no change between the percent of reads predicted to be derived from coding domain sequences when a species-level database is used versus when all of GTDB is used to predict open reading frames The slight increase observable for some species is a result of different thresholds, where we used 0.39 for the species database and 0.5 for the GTDB rs202 database.

**Table S1:**
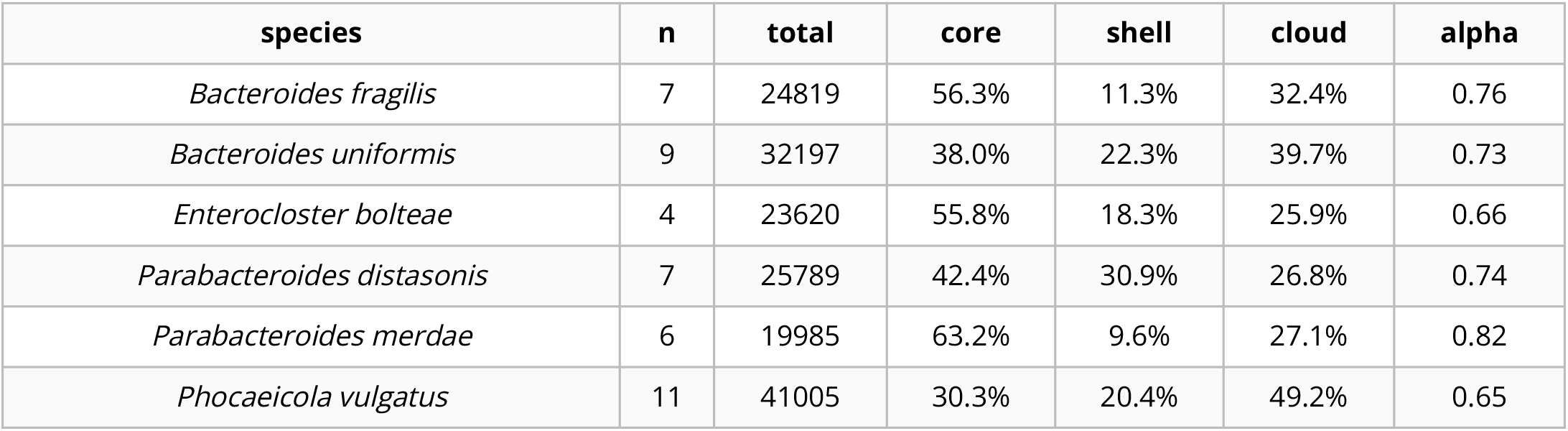
Metapangenome estimates for each species. *n* designates the number of metagenomes used to estimate the total, core, shell, cloud, and alpha values. Unlike isolate genomes, metagenomes may contain a fraction of an organism’s genome if the metagenome was not sequenced deeply or if an organism was rare. To calculate the core, shell, and cloud fractions and to estimate the openness of the metapangenome, we removed samples with fewer than 10,000 k_aa_-mers.

**Figure S4:**
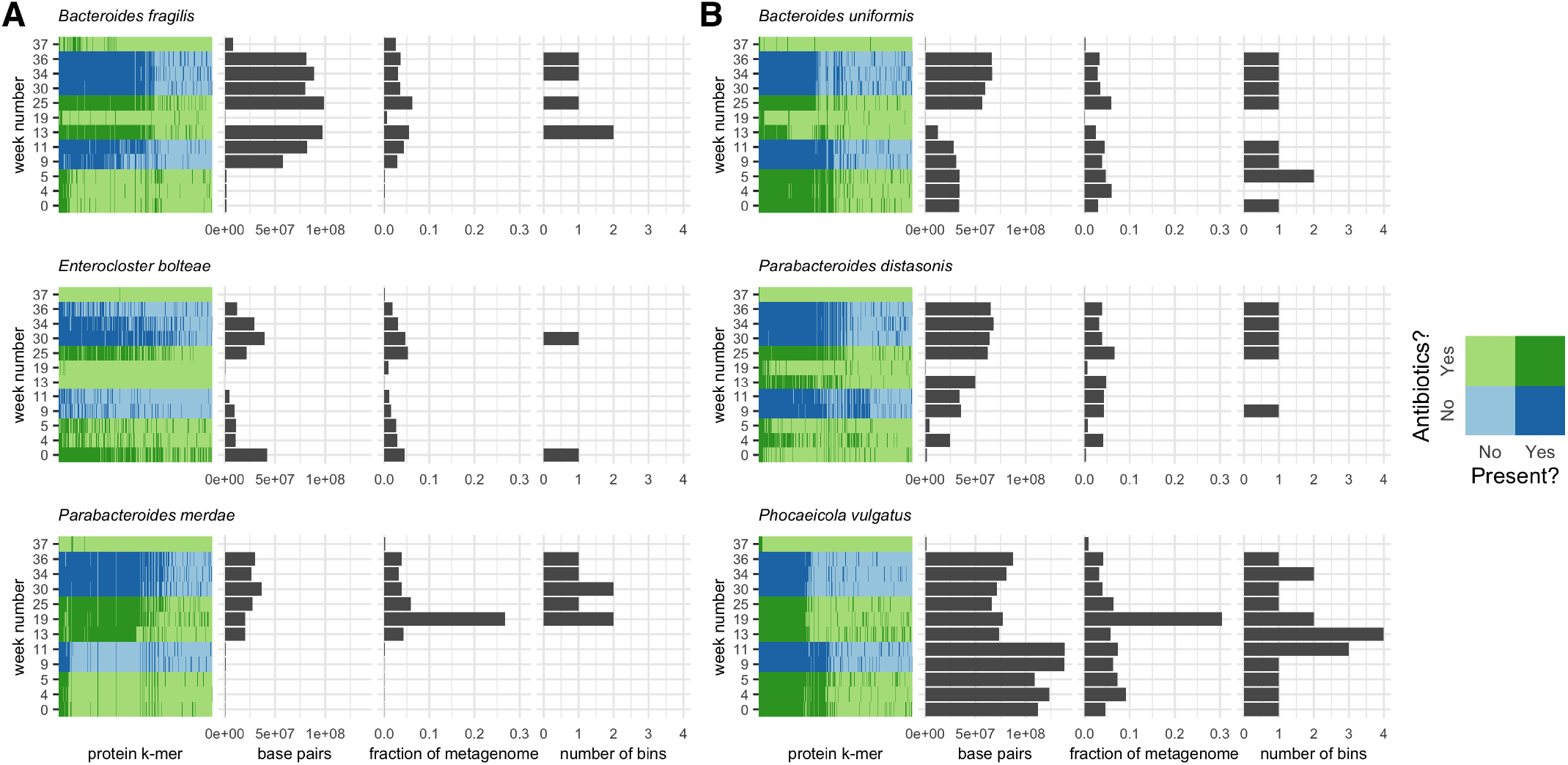
k_aa_-mer metapangenomes for six species. Each species contains a four-panel figure. The first panel is a binmap plot. Dark colors represent k-mers that are present in each sample. Blue shades represent time points when the sampled individual was not on antibiotics, while green shades represent time points when the individual was on antibiotics. The second panel represents an estimated number of base pairs in the metagenome detected to originate from that species. The third panel represents an estimated fraction of the metagenome assigned to that species. The fourth panel represents the number of bins produced for that species from that sample using a *de novo* metagenome assembly and binning approach. The two values represented in the second and third panels and the species assignations used to infer the value represented in the fourth panel were inferred using the sourmash gather algorithm against the GTDB rs202 database. **A)** Species for which presence-absence fluctuated over the time series. **B)** Species for which strain presence-absence fluctuated over the time series.

**Table S2:**
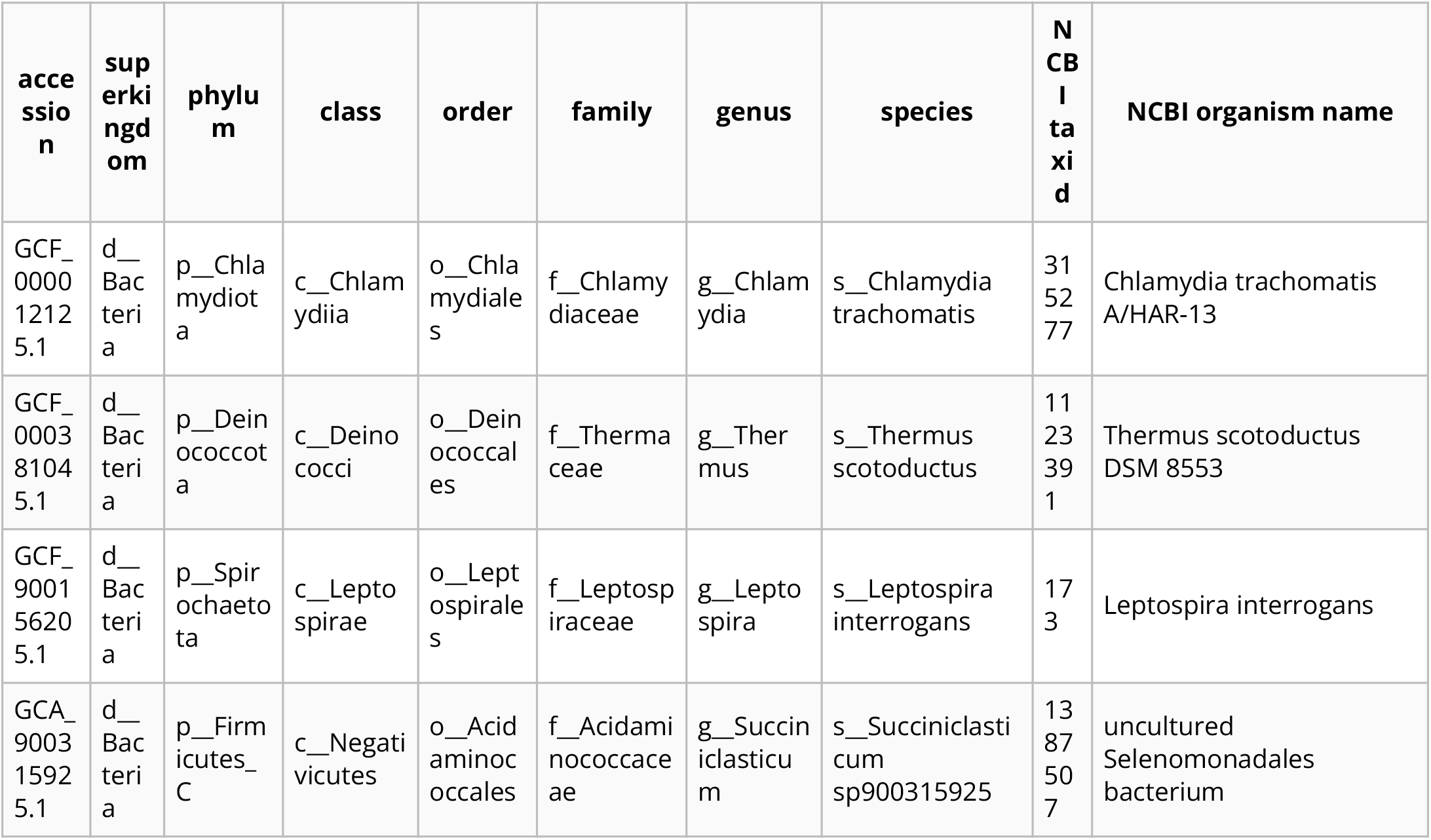
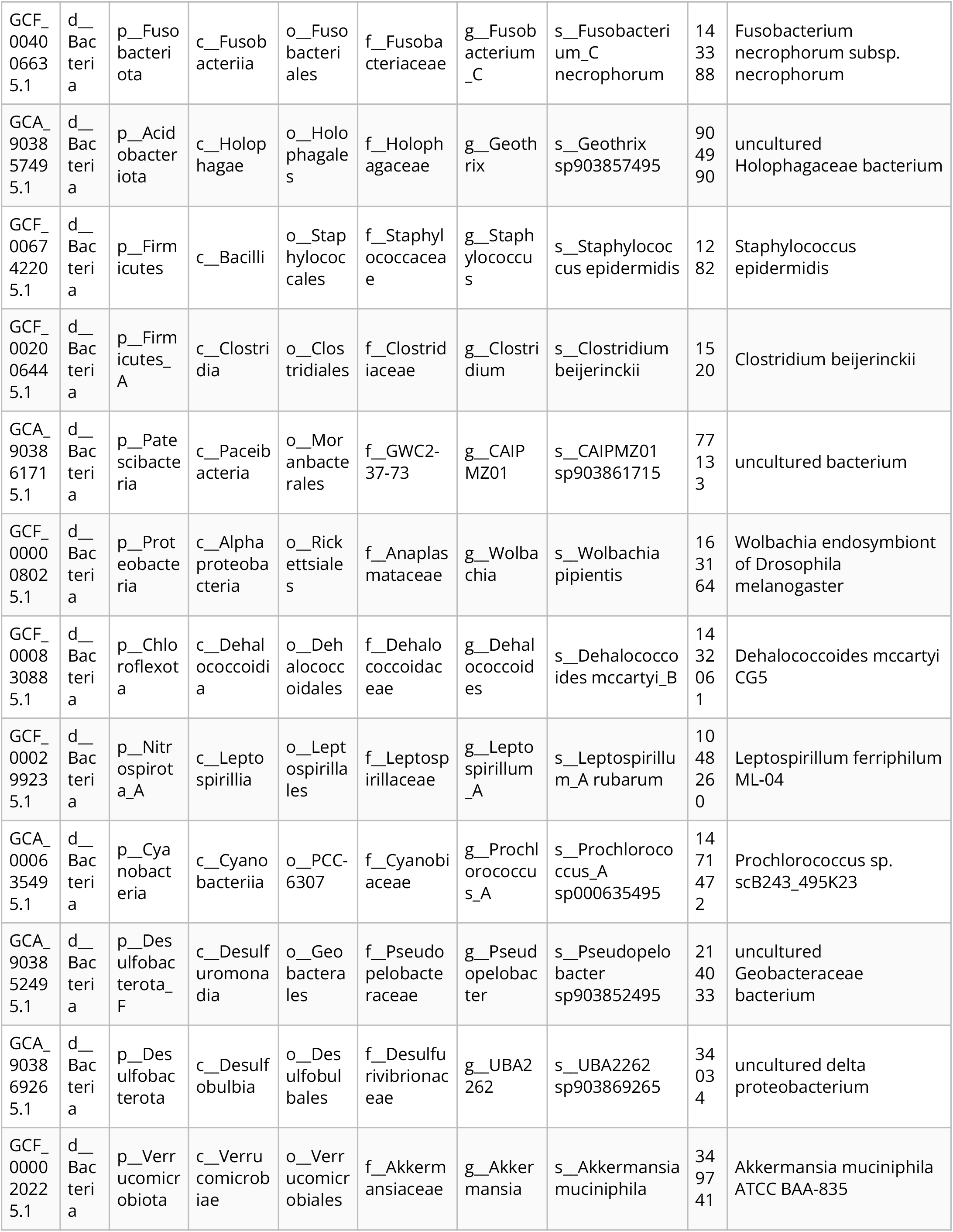
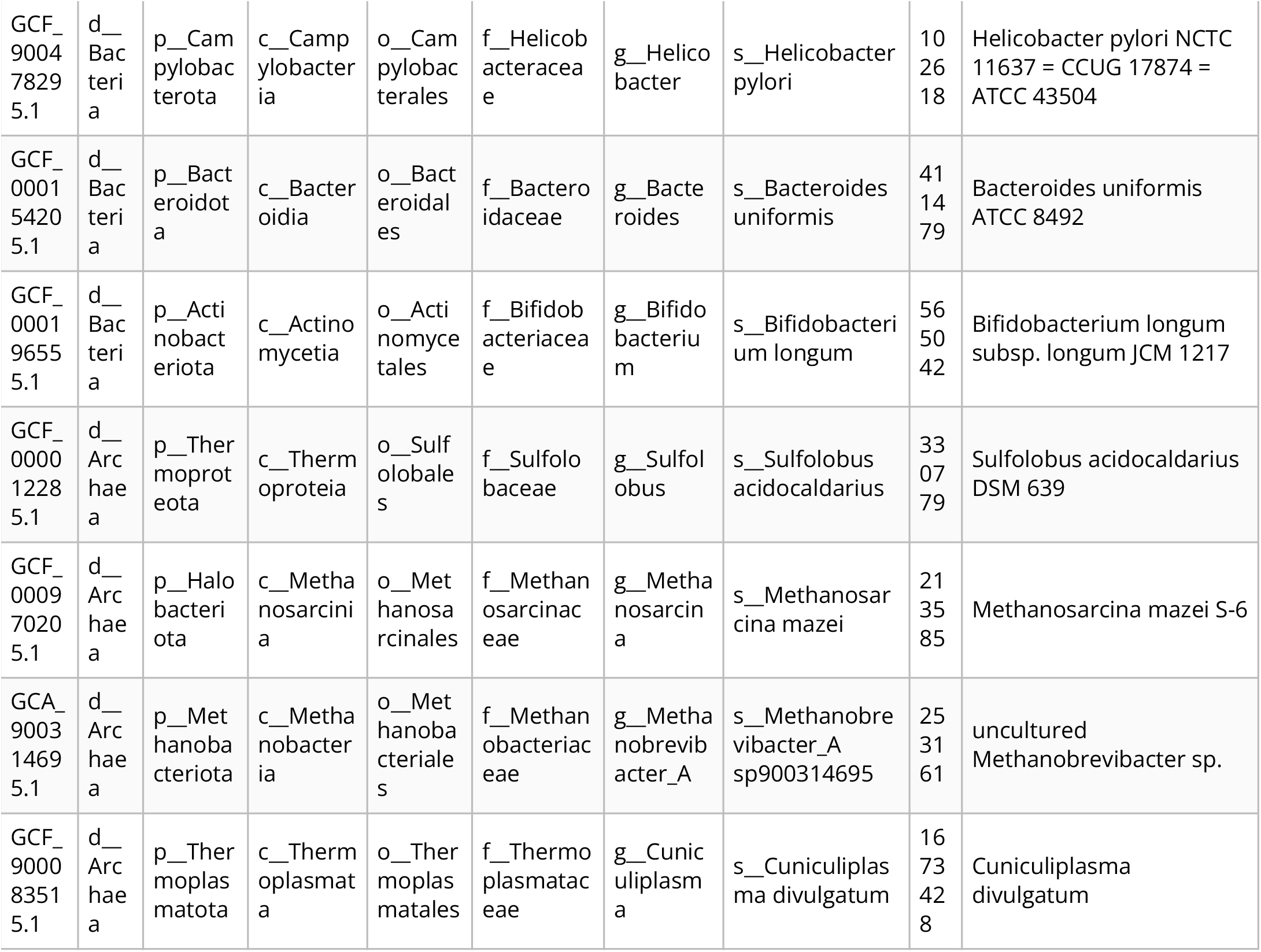
GTDB genomes used to benchmark orpheum accuracy.

**Table S3:**
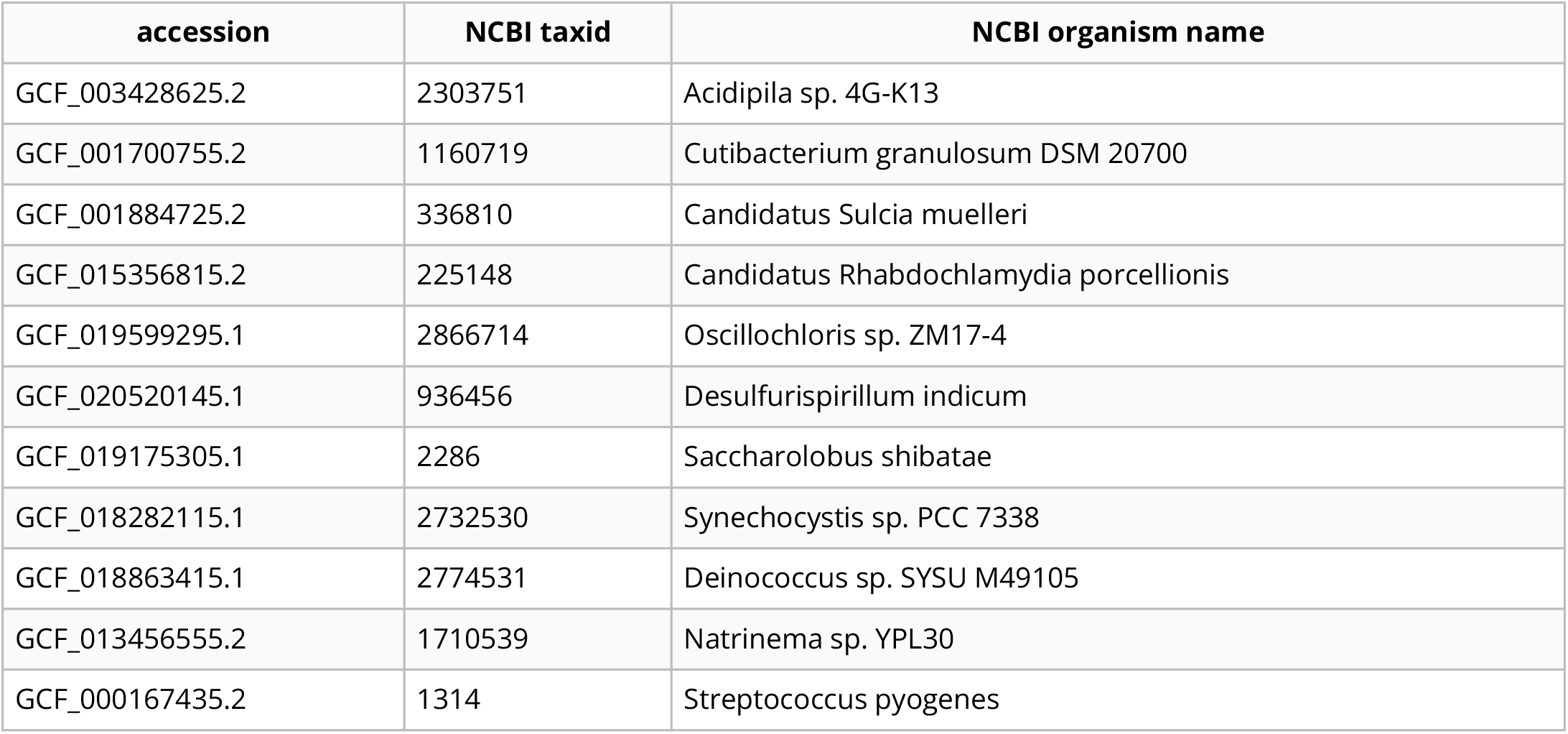
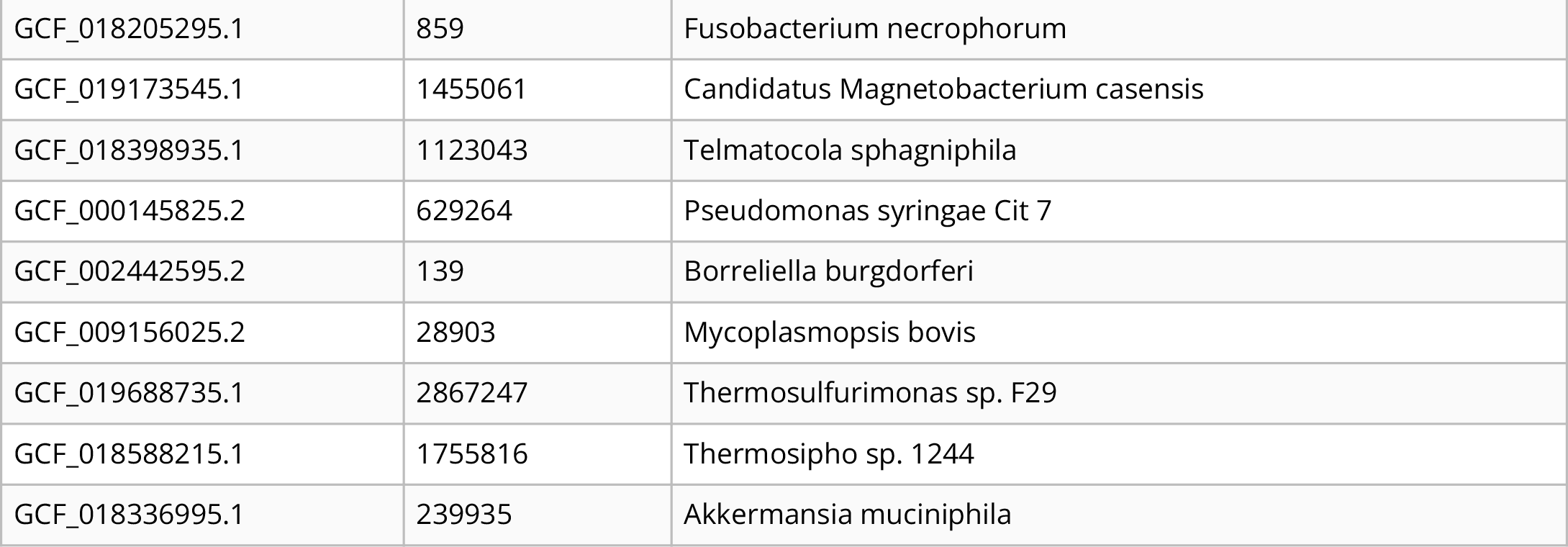
RefSeq genomes not in the GTDB rs202 database used to benchmark orpheum accuracy.

## Practical considerations for building k_aa_-mer metapangenomes

### Open reading frame prediction with orpheum

The RAM used to run orpheum is dictated by the database size, as the database is loaded into to memory while it is running. The GTDB rs202 nodegraph was 94 GB in size, and the RAM required to run orpheum never exceed 97GB, which makes database distribution and orpheum execution available on high performance compute clusters and other remote computers. Alternatively, species level databases were ∼5 Mb in size, reducing the RAM and CPU time needed to run orpheum.

We demonstrated that orpheum is better able to predict open reading frames in genomes that have species-level representatives in the GTDB database. To asses whether this criteria is satisfied by a query genome without performing genome assembly, we recommend sourmash gather [19]. Sourmash gather will estimate the fraction of sequencing reads in a genome or metagenome that match to genomes in GTDB by comparing long nucleotide k-mers in the query against those in the database [19]. Alternatively, the tool SingleM could be used to perform this task (https://github.com/wwood/singlem). SingleM estimates the taxonomic composition of sequencing reads by identifying fragments of single copy marker genes in short reads and comparing them against a database of taxonomically labelled sequences.

These strategies may also be useful to predetermine the set of species-level databases to use for ORF prediction.

### K_aa_-mer pangenome construction

One consideration is for k_aa_-mer sketch creation is the scaled value. The scaled parameter controls the fraction on k_aa_-mers included in each sketch. We have found that a scaled value of 100 works well for comparing proteomes [17]. For a subset of pangenomes benchmarked in results section, “K-mer methods accurately predict open reading frames in short sequencing reads”, we tested scaled = 1 and scaled = 100 (n = 8; s Akkermansia muciniphila, s Chlamydia trachomatis, s Faecalibacterium prausnitzii_D s Gemmiger formicilis, s Geothrix sp903857495, s Methanobrevibacter_A-sp900314695, s Thermus scotoductus, s UBA2262 sp903869265). We correlated the number of k_aa_-mers and genes per pangenome, the number of unique k_aa_-mers and genes per pangenome, and the similarity between genomes using k_aa_-mers or genes with sketches generated using a scaled of 1 or a scaled of 100. We then performed a second order correlation to determine if both scaled values produced the same results. All values were strongly significantly correlated with an R^2^ > 0.99 (p < 0.001), indicating that scaled 100 sketches captured the same patterns as scaled 1 sketches. While we have not experimented with the upper bound of the scaled value that will still produce accurate results, using a scaled of 100 substantially decreased compute times.

